# Age-dependent Powassan Virus Lethality is Directed by Glial Cell Activation and Divergent Neuroinflammatory Cytokine Responses in a Murine Model

**DOI:** 10.1101/2023.12.18.572230

**Authors:** Megan C. Mladinich, Grace E. Himmler, Jonas N. Conde, Elena E. Gorbunova, William R. Schutt, Shayan Sarkar, Stella E. Tsirka, Hwan Keun Kim, Erich R. Mackow

## Abstract

Powassan virus (POWV) is an emergent tick-borne flavivirus that causes fatal encephalitis in the elderly and long-term neurologic sequelae in survivors. How age contributes to severe POWV encephalitis remains an enigma and there are currently no animal models that reflect age-dependent POWV neuropathology. Inoculating C57BL/6 mice with a POWV strain (LI9) currently circulating in *Ixodes* ticks, resulted in age-dependent POWV lethality with overt spongiform brain damage 10-15 dpi. Infection of 50 week old mice resulted in 82% lethality 10-15 dpi that was sequentially reduced by age to 7.1% in 10 week old mice. LI9 encephalitis resulted in early neuronal depletion, with severe CNS damage, persistent inflammatory gliosis and long-term spongiform pathology in survivors (30 dpi). In all mice POWV LI9 was neuroinvasive and reached maximum POWV loads in the CNS 10 dpi. Coincident with murine lethality, in 50 week old mice maximum POWV CNS levels persisted 15 dpi, while instead decreasing by 2-4 logs in 10-30 week old mice. Although glial cells were highly activated in all POWV infected mice, differences in age-dependent CNS cytokine responses were striking 15 dpi. In 50 week old mice POWV induced Th1-type cytokines (IFNγ, IL-2, IL-12, IL-4, TNFα, IL-6), suggesting a pro-inflammatory M1 microglial activation cascade. In contrast, POWV induced Th2-type cytokines (IL-10, TGFβ, IL-4) in 10 week old mice consistent with a neuroprotective M2 microglial phenotype. These findings reflect differences in neurodegenerative versus neuroprotective glial cell responses that correlate with divergent CNS viral clearance and age-dependent POWV LI9 lethality. Discrete age-dependent CNS cytokine responses suggest neuroinflammatory targets as potential POWV therapeutics. These studies establish a highly lethal POWV murine model and reveal a hyperinflammatory mechanism of age-dependent POWV lethality that mirrors human POWV severity and long-term CNS sequelae in the elderly.

**Importance:** Powassan virus is an emerging tick-borne flavivirus causing lethal encephalitis in aged individuals. We reveal an age-dependent POWV murine model that mirrors human POWV encephalitis and long-term CNS damage in the elderly. Findings demonstrate that POWV load and discrete glial cell cytokine responses in the CNS are critical determinants of age-dependent POWV lethality. POWV age-independently activates microglia and astrocytes, but directs neuroprotective Th2 cytokine responses in 10 week old mice and distinct pro-inflammatory Th1 cytokine responses in the CNS of 50 week old mice. This reveals roles for a hyperinflammatory CNS cytokine cascade in age-dependent POWV lethality, and protective anti-inflammatory cytokines in murine survival. Notably, results define potential therapeutic targets and rationalize approaches for preventing severe POWV encephalitis that may be broadly applicable to neurodegenerative diseases. This age-dependent murine POWV model permits analysis of vaccines, and therapeutics that prevent POWV neuroinvasion or resolve severe POWV encephalitis in the elderly.

## INTRODUCTION

Flaviviruses (FVs) are a family of enveloped, positive-strand RNA viruses that cause human disease, and numerous FVs are transmitted by arthropod vectors(^1^). Tick-borne FVs cause ∼15,000 annual cases of severe encephalitis and include the tick-borne encephalitis virus (TBEV), found in Eurasia, and Powassan virus (POWV), in North America(^2^). POWV (strain LB) was first isolated in 1958 from a human encephalitis case in Powassan Ontario Canada(^3^). POWVs are emerging in the Northeastern United States due to the expansion of animal reservoirs, tick vectors and an increasing clinical awareness of human POWV infections(^2, 4–13^). In endemic US states, the seroprevalence of POWV is between 0.7-6.1%, with fewer clinical cases suggesting that infections are largely asymptomatic(^2, 14^). POWV is present in tick saliva, transmitted in as little as 15 minutes of a tick bite(^2, 11, 15^), and in severe cases causes 10% fatal encephalitis and long-term neurologic sequalae in 50% of survivors(^2, 16^). POWV infects all age groups, with limited data suggesting that lethality and severe neurologic sequelae are increased in patients >60 years of age, similar to clinical outcomes of TBEV infection(^2, 13, 17, 18^). POWV infections are biphasic with an initial acute febrile illness 1-3 weeks after a tick bite followed by weeks of wide-ranging CNS manifestations(^2, 16, 17^). Fatal human POWV encephalitis presents with severe CNS damage, brainstem involvement, inflammation and gliosis in the cerebral cortex, without histologic evidence of systemic infection(^2, 16, 17^). Currently, there are no clinically approved POWV therapeutics or vaccines.

There are 2 POWV genotypes (Lineage I and II) that reflect their primary animal and tick hosts(^19, 20^). However, with 96% amino acid identity in their Envelope (Env) proteins(^21, 22^), POWV strains are serologically indistinguishable and comprise a single serotype, suggesting the broad efficacy of a POWV vaccine(^23^). Animal models are critical for analyzing vaccines, therapeutics and resolving mechanisms of viral neuropathogenesis. Prior POWV murine infection models utilized neuroadapted POWV strains (LB, SP, IPS1) first isolated and serially passaged in murine brains(^2, 3, 10, 12, 22, 24–29^). Infection of 5-14 week old mice with POWV LB results in rapid lethality 6-9 dpi in >60% of mice, while infection with POWVs SP or IPS1 results in delayed (9-10 dpi) or varied mortality(^2, 12, 27, 28^). Aside from CNS involvement, there is little understanding of POWV directed encephalitis or the mechanism of POWV lethality in mice(^27, 29–33^). Lethality has been reported to vary by strain, inoculation titers and sites, the involvement of tick salivary glands, and the timing of POWV neuroinvasion. Further, lethality may be impacted by the use of strains neuroadapted to replicate in murine brains, and the use of young 6-14 week old mice(^10, 24, 25, 27–29, 34–36^).

TBEV encephalitis is characterized by neuronal damage and glial cell involvement(^2, 37–39^). CNS pathogenesis includes vacuolation of the neuropil, widespread inflammatory CNS infiltrates, sparse infected Purkinje cells, and dispersed large neurons in the medulla, pons, cerebellum and striatum(^39, 40^). However, severe inflammation was associated with reduced TBEV antigen and in a third of human cases TBEV antigen was undetectable in the CNS(^39, 40^). In a large TBEV case study, infection was reported in all age groups, with no fatal cases in patients <40 years of age, and increasing fatality rates in patients 60-90 years old(^2, 37–39^).

In comparison to TBEV, the CDC reported that 8% of POWV patients were <18 years old and ∼50% of patients were >60 years old(^2^). In a cohort of 99 POWV cases, the median age was 62 years and all fatalities (11%) occurred in patients older than 50 years of age(^7^). In New York, analysis of 14 NY encephalitis cases revealed that 72% of patients were >60 years old, that all 5 fatalities were in patients >60 years of age and that all survivors had neurologic deficits(^14, 17^). Autopsies revealed reactive gliosis, increased microglia and necrotizing CNS inflammation consistent with acute meningoencephalitis(^2, 14^). These findings suggest age as a factor in severity of encephalitis caused by TBEV and POWV.

In a 2020 survey, 2% of *I. scapularis* ticks in Long Island, NY were POWV positive, and POWV strain LI9 was isolated in Vero cells directly from *Ixodes* ticks, without murine neuroadaptation(^21, 41^). POWV LI9 nonlytically infects VeroE6 cells, spreading cell-to-cell in the presence of neutralizing antibodies(^21, 42^), and *in vitro*, infects primary human brain microvascular endothelial cells (hBMECs) that form the blood-brain-barrier(^21^). Basolateral spread from polarized hBMECs suggests a mechanism for POWVs to enter neuronal compartments(^21^), although *in vivo* CNS entry mechanisms for POWV have not been assessed. C57BL/6 mice subcutaneously inoculated with POWV LI9 seroconvert and produce cross-reactive POWV neutralizing antibodies(^21, 42^).

Here we evaluated POWV LI9 neurovirulence and lethality as a function of age across 10-50 week old C57BL/6 mice. Footpad inoculation of POWV LI9 resulted in 82% lethality in 50 week old mice 10-15 dpi, with lethality reduced sequentially with age to 7.1% in 10 week old mice. Despite age-dependent differences in lethality, all infected mice elicited POWV neutralizing antibodies, and harbored POWV LI9 RNA in the CNS. Neuronal depletion and spongiform damage was present by 5 dpi, and spongiform lesions were present throughout LI9 infected brains with pronounced damage in the cerebellum, hindbrain and brainstem. CNS pathology revealed neuronal necrosis, spongiform encephalopathy, and gliosis, 5-15 dpi(^43–49^). In 10-40 week old mice, CNS viral loads increased to maximal levels 10 dpi. By 15 dpi, CNS POWV RNA levels were reduced by 2-3 logs in 10-40 week old mice, but remained at high levels in 50 week old mice. Spongiform encephalopathy and microgliosis persisted in the CNS of surviving mice 30 dpi, consistent with POWV causing long-term neurologic damage in the elderly.

Analysis of CNS responses to POWV in 50 week old mice revealed the robust induction of pro-inflammatory Th1-type cytokines (IL-2, IL-6, IL-12, IL-1β, TNFα, IFNγ) 15 dpi, consistent with a neurodegenerative M1 microglial phenotype(^46, 48, 50–59^). Contrastingly, in 10 week old mice POWV induced Th2-type cytokines (IL-10, TGF-β, IL-4) and chemokines in the CNS that are consistent with a neuroprotective M2 microglial phenotype(^53, 57, 60, 61^). This advances a mechanism of POWV lethality determined by opposing age-dependent glial cell activation phenotypes that either promote a neurodegenerative inflammatory cascade, or direct neuroprotective responses, which govern viral clearance from the CNS(^43, 48, 58, 59, 62, 63^).

Studies establish an age-dependent murine model of POWV neurovirulence that reflects severe lethal disease and persistent cognitive impairment in the elderly (^2, 7, 13, 14^). Findings reveal hyperinflammatory CNS cytokine responses and limited POWV CNS clearance as determinants of age-dependent POWV lethality, and suggest potential therapeutic targets and approaches for preventing lethal POWV disease.

## RESULTS

### POWV LI9 Lethality is Age-Dependent in C57BL/6 Mice

POWV LI9 is a currently circulating virus isolated directly from *Ixodes* ticks in VeroE6 cells(^21^). Tick transmitted TBEV and POWV are associated with the age-dependent severity of human encephalitis(^2, 7, 14, 17, 37–39^). To investigate age as a determinant of LI9 lethality, we footpad inoculated 10, 20, 30, 40 and 50 week old C57BL/6 mice with LI9 (2 × 10^3^ FFU) and kinetically assessed lethal neurovirulent disease (Figure 1). In all age groups, LI9 infection resulted in weight loss and clinical neurologic symptoms including hindlimb flaccid paralysis, ataxia and weak grip 8-15 dpi. In 10 week old mice, initial weight loss 8-10 dpi was fully recovered in surviving mice by 15 dpi, while 20-50 week old mice only partially recover body weight 15-20 dpi (Figure 1A). POWV LI9 lethality was primarily observed 10-14 dpi (Figure 1B) with a sequential age-dependent increase in lethality from 7.1% (10 week), 20% (20 week), 30% (30 week), 39% (40 week) to 82% (50 week), (n=11-20/group; Figure 1B,E). All mice seroconverted and produced neutralizing POWV antibodies as determined by focus neutralization assay (IC50 range: 7 × 10^2^ to 1 × 10^4^)(Table 1). Footpad inoculation of 50 week old mice with 2 or 20 FFUs of LI9 or PBS (n=20) resulted in neurologic symptoms, weight loss 8-17 dpi (Figure 1C) and lethality in 45% versus 80% of mice, respectively (Figure 1D, E). The mortality of LI9 infected male and female mice, summed across all ages or specifically in 50 week old mice, was not statistically significant (Figure 1E,F). Collectively this data reveals that a minimal infectious dose of POWV LI9 (2-20 FFU) is highly lethal in 50 week old mice and that POWV LI9 lethality increases with the age of C57BL/6 mice.

**Figure 1.**
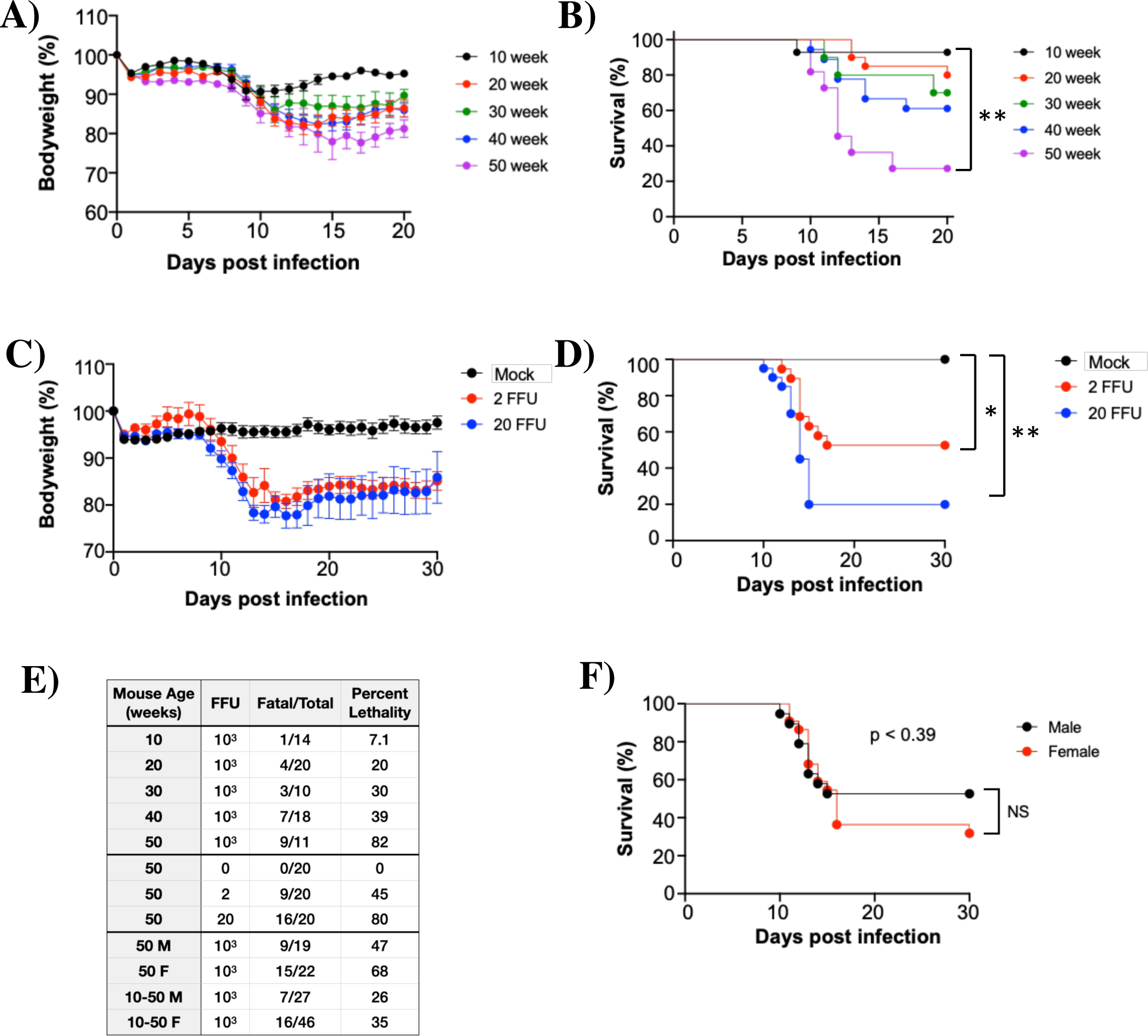
The Lethality of POWV LI9 is Age-dependent in C57BL/6 Mice. (A-F) C57BL/6 mice 10-50 weeks old were footpad inoculated with 2 × 10^3^ FFU (A, B, F), or 2-20 FFU (C, D) of POWV LI9 or PBS. (A, C) Mice were weighed daily, with body weight of surviving mice expressed as a percent of weight prior to infection (mean +/- standard error of the indicated number of mice per group). (B, D) Kaplan-Meier curves of lethal POWV infection of 10-50 week old mice (B): 10 wk (n=14), 20 wk (n=20), 30 wk (n=10), 40 wk (n=18), 50 wk (n=11) old mice; and (D) 50 week old mice inoculated with 2 FFU (n=20), 20 FFU (n=20), or mock infected (n=20) were analyzed by a log rank test (\**p<*0.05; \*\**p<*0.01). (E) Summary of murine age, POWV LI9 inoculation dose, fatal/total infected and percent lethality. Overall lethality of POWV in male (n=27) vs. female (n=46) mice across all ages was not significant (NS; p<0.41). (F) Kaplan-Meier curve of lethal POWV infection of 50 week old male (n=19) versus 50 week old female (n=22) mice inoculated with 2 × 10^3^ FFU, analyzed by a log rank test, were not significant (NS; p<0.39).

**Table 1.** POWV Infected Mice Seroconvert to Produce LI9 Neutralizing Antibodies. C57BL/6 10-50 week old mice were s.c. footpad inoculated with 2×10^3^ FFU of POWV LI9 or mock-infected with PBS. Neutralizing antibodies present in POWV LI9 or mock-infected mouse serum 30 dpi were assessed using a POWV LI9 focus reduction neutralization assay in VeroE6 cells. Serial dilutions of sera were added to 500 FFU of POWV LI9 prior to adsorption to cells and 36 hpi infected VeroE6 cells were immunostained using anti-POWV-HMAF and quantitated to define IC50s. Serum dilutions required to reduce POWV foci by 50% are presented as a range (n=4/age group).

| Mouse Age<br>(weeks) | Neutralizing Ab IC50<br>(30 dpi) |
| --- | --- |
| 10 | $1.0 \times 10^3 - 2.0 \times 10^3$ |
| 20 | $1.2 \times 10^3 - 2.7 \times 10^3$ |
| 30 | $9.8 \times 10^2 - 2.4 \times 10^3$ |
| 40 | $6.8 \times 10^2 - 1.6 \times 10^3$ |
| 50 | $4.6 \times 10^3 - 1.2 \times 10^4$ |

### POWV Causes Spongiform Encephalitis and CNS Resident Glial Cell Activation

Pathology and CNS inflammation associated with lethal POWV LI9 neurovirulence were evaluated on formalin-fixed brains from mock and LI9 infected 10 and 50 week old mice, 5 and 15 dpi. Representative brain sections were assessed for histopathology by H&E staining, and for activated microglia/macrophages by Iba1 immunostaining. H&E staining of the POWV infected pons in 10 and 50 week old mice revealed rapid spongiform encephalopathy and neuronal necrosis by 5 dpi that was also apparent 15 dpi, and absent in mock infected 50 week old controls (Figure 2A). Increased microgliosis, with the appearance of Iba1^+^ microglial nodules(^64, 65^), was observed 5 dpi and amplified 15 dpi in POWV infected 10 and 50 week old mice versus controls (Figure 2A). Scoring of the pons 5-15 dpi for spongiform encephalopathy, microgliosis and neuronal necrosis (n=4) revealed similar pathology increases in 10 and 50 week LI9 infected mice versus controls (Figure 2B). ImageJ quantification of Iba1^+^ staining (10 areas per mouse, n=3) showed significant increases in the pons of both 10 and 50 week old POWV infected brains 15 dpi versus mock infected controls (Figure 2C), and reflect an increase in microglia/macrophages in the POWV infected CNS. POWV directed neuronal loss was confirmed by decreased immunostaining of neurons (NeuN antibody) at both 5 and 15 dpi versus mock infected controls (Figure S1A). Purkinje cells are reported encephalitis targets of TBEV, POWV LB and West Nile virus (WNV)(^29, 33, 39, 66, 67^), and noted at autopsy of immunosuppressed POWV patients(^16, 68^). Despite the spongiform CNS appearance of the POWV LI9 infected CNS, we found no histologic evidence of infection, depletion or damage to granular, Purkinje or molecular layers of the cerebellum in 10 or 50 week old mice (5, 15 or 30 dpi) versus mock infected 50 week old controls (Figure S1B).

**Figure 2.**
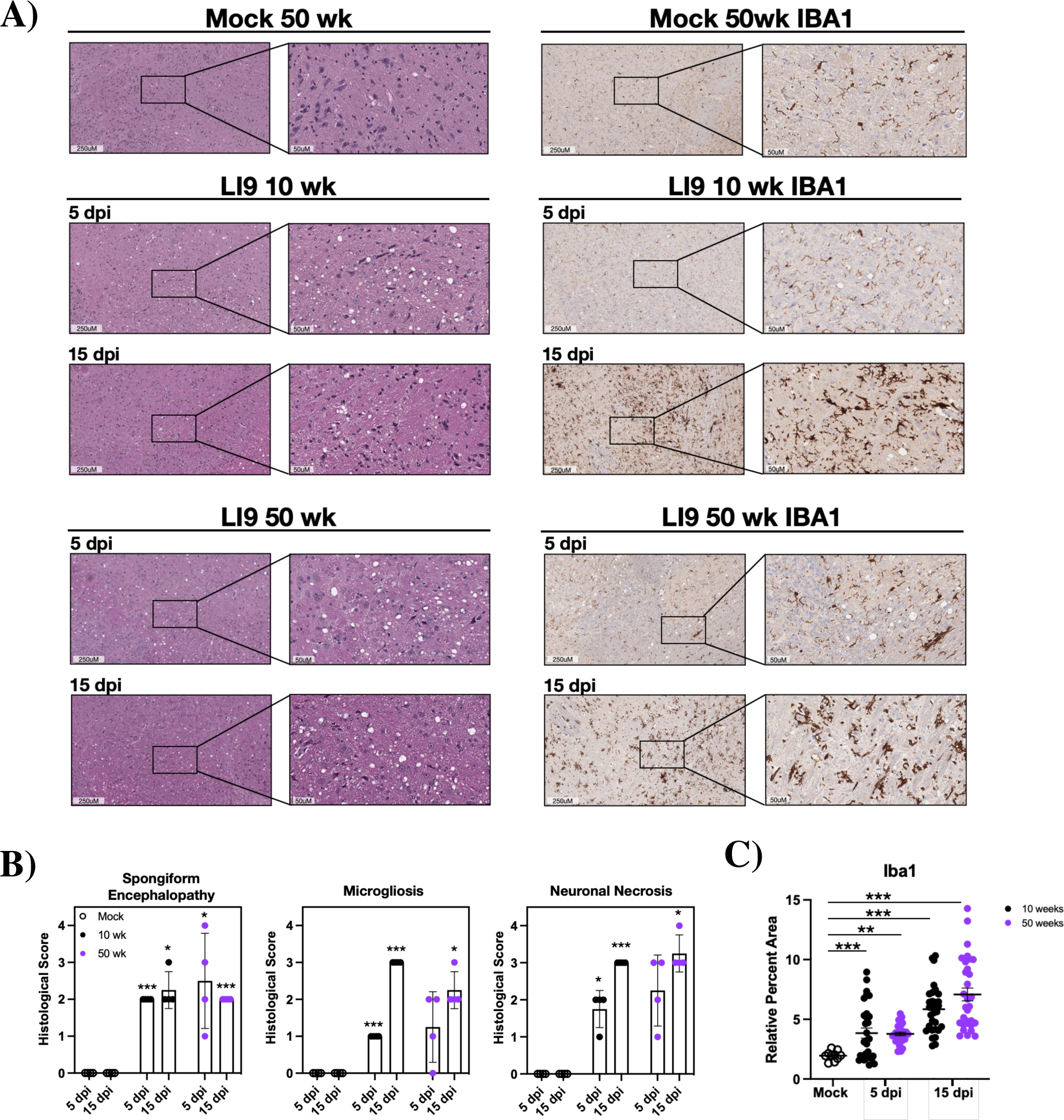
Kinetic Changes in the Pons of POWV Infected Mice. (A) C57BL/6 10 and 50 week old mice were footpad inoculated with 2 × 10^3^ FFU of POWV LI9 or mock infected with PBS. Brains were harvested 5 and 15 dpi, and H&E stained (n=4), or immunostained for microglia/macrophages (n=3) using Iba1 antibody. Representative sections of the Pons are presented. (B) The severity of POWV directed spongiform encephalopathy, microgliosis, and neuronal necrosis in H&E brain regions (n=4) was scored on a scale of 0-4 by blinded comparison versus age-matched controls: (0) baseline determined by control brain staining in select region, (1) localized lesion, (2) multiple localized lesions, (3) lesions spread throughout most of select region, (4) lesions uniformly spread throughout select region. Two-way comparisons were performed by two-tailed analysis of variance and an unpaired Student’s t-test. (C) ImageJ quantification of Iba1 immunostaining: relative percent area of Iba1^+^ pixel intensity was determined for 10 CNS regions of 10 and 50 week old mice (n=3), 5 and 15 dpi, vs an age-matched mock infected controls. Comparisons to mock infected controls were performed by two-way ANOVA analysis. Asterisks indicate statistical significance (*p*<*0.01; **p*<*0.001, ***p*<*0.0001).

We comprehensively analyzed brain regions of POWV infected 10 and 50 week old mice 15 dpi. We observed striking spongiform encephalopathy with overt H&E stained histologic lesions, severe neuronal necrosis, glial scarring and perivascular cuffing in the pons, medulla, brainstem, and cerebellum compared to age-matched controls (Figure 3A)(^69^). Increases in inflammatory Iba1^+^ staining of microglia/macrophages was apparent in all CNS regions in both 10 and 50 week old mice (Figure 3A). Comparative scoring of CNS regions of both 10 and 50 week old mice reflected vastly increased spongiform encephalopathy and severe microgliosis throughout POWV infected brain regions (Figure 3B). Similar H&E and Iba1 staining, spongiform damage, microgliosis and neuronal necrosis were observed in the CNS of 20, 30 and 40 week old mice 5 and 15 dpi (Figure S2A) with similar scoring across murine ages (Figure S2B) and CNS regions (Figure S3).

**Figure 3.**
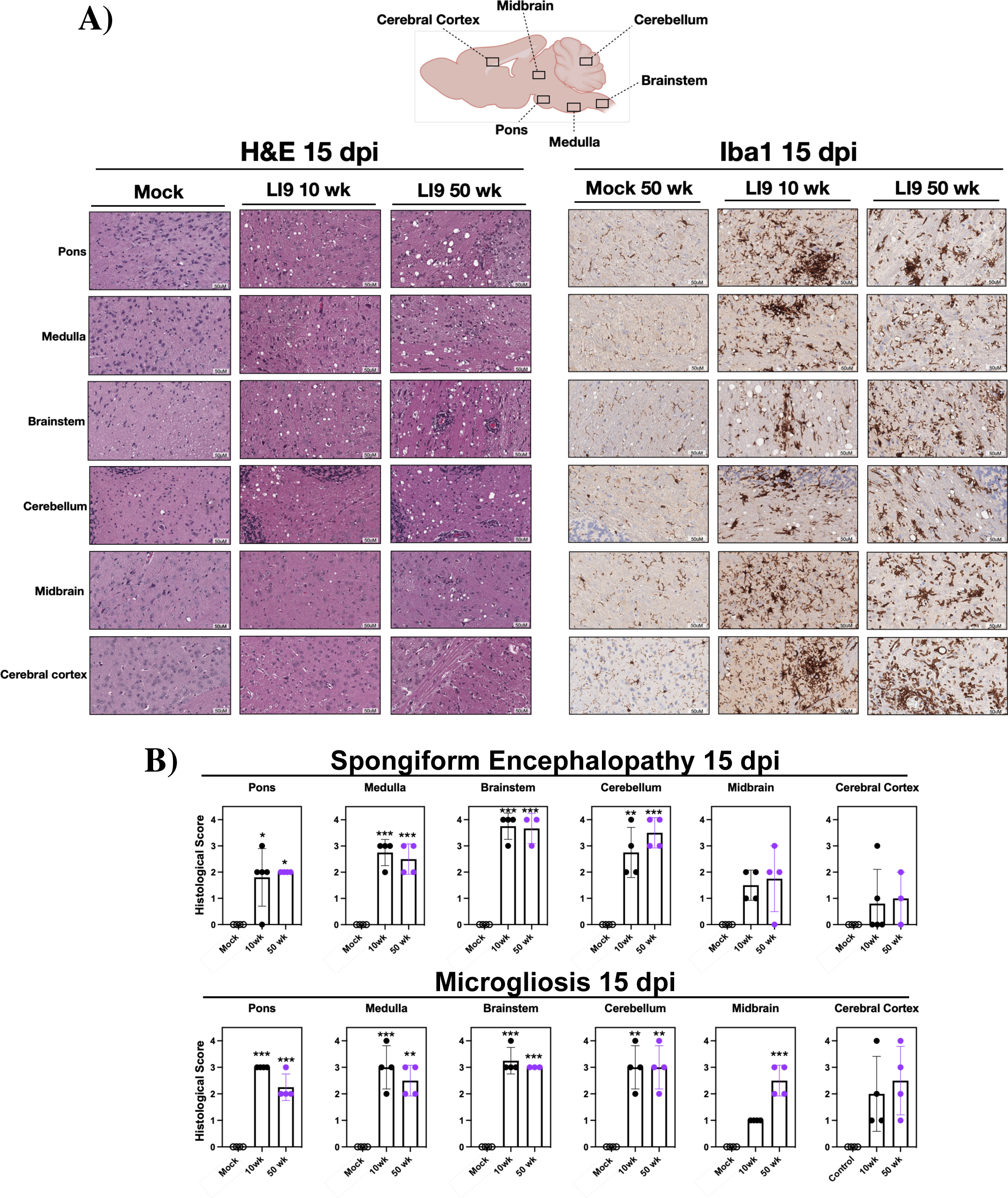
POWV Causes Spongiform Encephalitis and Microgliosis in Mice 15 dpi. (A, B) 10 and 50 week old C57BL/6 mice were footpad inoculated with 2 × 10^3^ FFU of POWV LI9 or mock infected with PBS. Brains were harvested 15 dpi, sectioned and either H&E stained (n=4) or immunostained for microglial/macrophage markers using anti-Iba1 (n=3). (A) Representative images of CNS histopathology in the Pons, Medulla, Brainstem, Cerebellum, Midbrain and Cerebral cortex of POWV versus mock infected 50 week old brains 15 dpi are presented. (B) The severity of POWV directed spongiform encephalopathy and microgliosis in H&E brain regions (n=4) was scored as in Figure 2 and compared to mock infected controls by two-tailed analysis of variance and an unpaired Student’s t-test. Asterisks indicate statistical significance (*p*<*0.01; **p*<*0.001, ***p*<*0.0001).

### CNS Damage and Microgliosis Persist in POWV Infected Survivors

POWV causes long term cognitive deficits in 50% of surviving patients(^2, 7, 14^). Analysis of brains from 10 and 50 week old mice that survived POWV infection (30 dpi) revealed persistent, long-term CNS spongiform pathology, increased microglial/macrophage staining (Iba1^+^) and neuronal necrosis versus age-matched controls (Figure 4A-C). Scoring of the Iba1 immunostained CNS 30 dpi demonstrated a significant increase in microglia/macrophages in 50 versus 10 week old POWV survivors and mock infected controls (Figure 4B). These results demonstrate that POWV is neuroinvasive and causes rapid CNS neuronal damage by 5 dpi with apparent increases in microglial/macrophage neuroinflammation from 5 to 15 dpi, that was maintained 30 dpi, in the CNS of both 10 and 50 week old mice.

**Figure 4.**
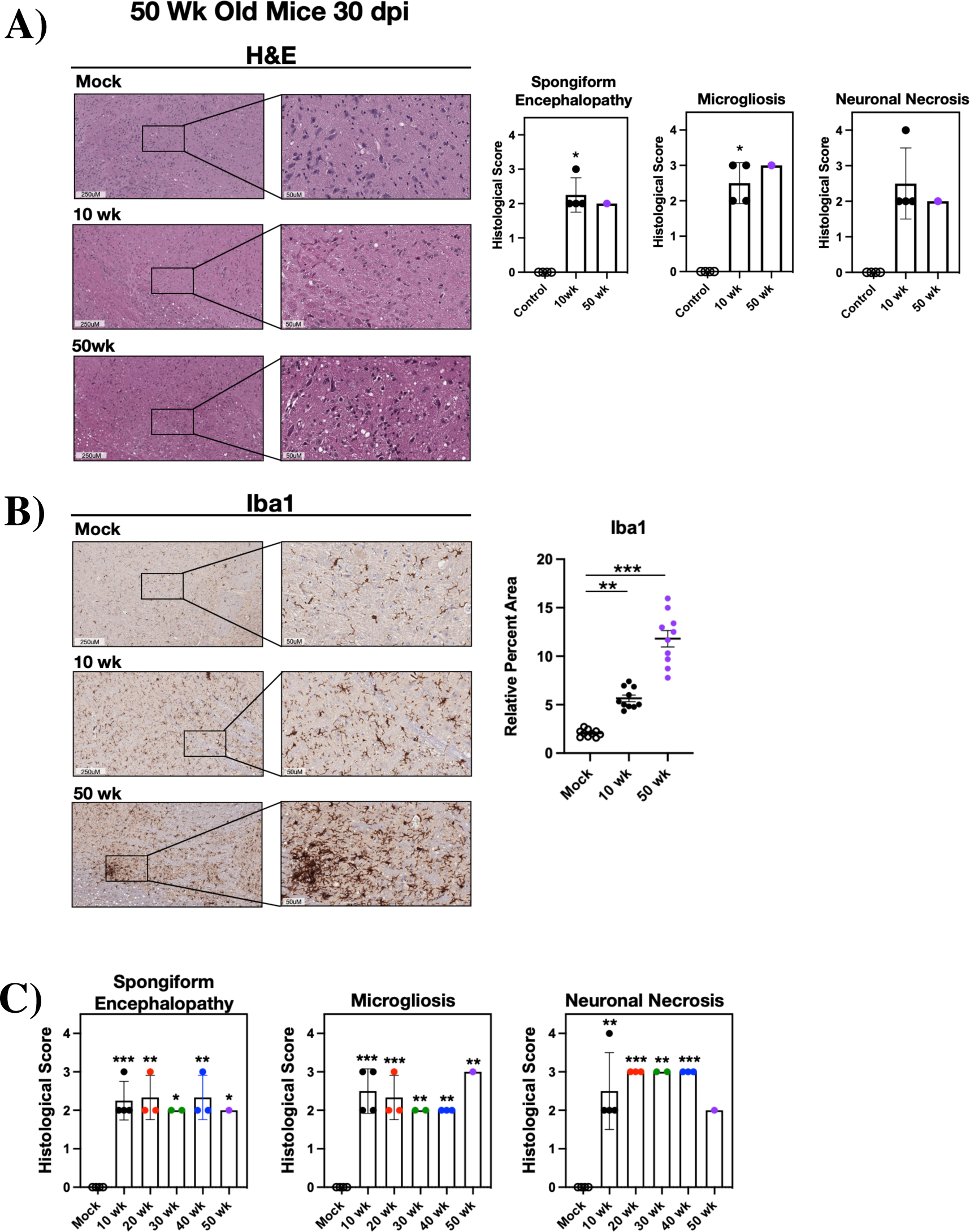
POWV Causes Persistent Spongiform Encephalopathy and Microgliosis. (A-C) 10 and 50 week old C57BL/6 mice that survived infection with 2 × 10^3^ FFU POWV LI9 were euthanized 30 dpi. Brains were harvested, sectioned and either (A) H&E stained (n=4) or (B) immunostained for microglial/macrophage markers using anti-Iba1 (n=3). (A) Representative images of CNS histopathology in the Pons of POWV versus mock infected 50 week old brains 30 dpi are presented (n=3). The severity of POWV directed spongiform lesions, microgliosis and neuronal necrosis was scored on a scale of 0-4 versus age-matched mock infected controls as in Figure 2. Two-way comparisons to mock infected age-matched controls were performed by two-tailed analysis of variance and an unpaired Student’s t-test. (B) Representative images of the Iba1 immunostained Pons of POWV infected versus mock infected 50 week old mice, 30 dpi (n=3) were quantitated by ImageJ on 10 CNS sections per mouse as in Figure 2C and compared to mock infected controls by one-way ANOVA analysis. (C) The severity of POWV directed spongiform encephalopathy, microgliosis, and neuronal necrosis in H&E Pons regions (n=4) was scored and analyzed, by two-way ANOVA, as in Figure 2. Asterisks indicate statistical significance (*p*<*0.01; **p*<*0.001, ***p*<*0.0001).

### POWV Infection Ubiquitously Activates Astrocytes In 10 and 50 Week old Mice

Astrocytes and microglia comprise CNS resident glial cells which direct immune surveillance and in combination amplify neuroinflammatory cytokine responses that orchestrate neurodegenerative and neuroprotective CNS responses(^44, 47–49, 58, 59, 62, 63^). To assess POWV activation of CNS astrocytes, we immunostained brain sections for expressed GFAP (glial fibrillary acidic protein)(^48, 70^). We found the GFAP was highly expressed in the POWV infected CNS, 5 and 15 dpi (Figure 5A), and across brain regions of both 10 and 50 week old mice (Figure 5B). Collectively our results demonstrate the activation of glial cells in the CNS of POWV infected mice independent of age.

**Figure 5.**
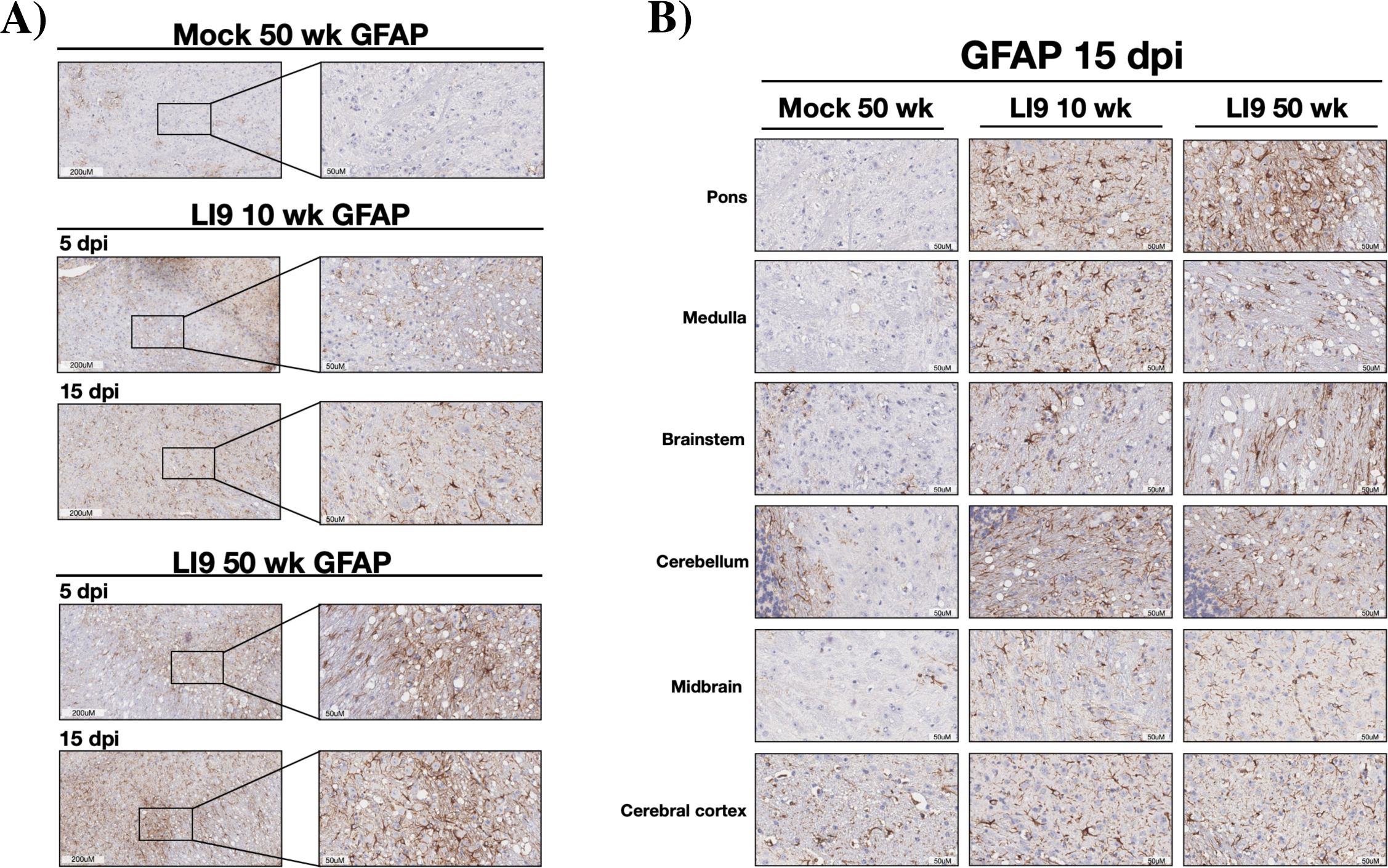
POWV Infection Causes Astrocyte Activation in Mice. (A-B) C57BL/6 10 and 50 week old mice were footpad inoculated with 2 × 10^3^ FFU of POWV LI9 or mock infected with PBS. Brains were harvested 5 or 15 dpi, sectioned, and immunostained for GFAP expressed in activated astrocytes. (A) Representative images of the GFAP immunostained Pons of POWV LI9 infected (n=1) and mock infected 50 week old (n=1) mice at 5 and 15 dpi. (B) Representative images of GFAP histopathology in the Pons, Medulla, Brainstem, Cerebellum, Midbrain and Cerebral cortex of 10 and 50 week old POWV infected brains versus mock infected 50 week old brains, 15 dpi, are presented.

### POWV Infected CNS Contain Few CD4 and CD8 T cell Infiltrates

While T cells in the POWV infected CNS have not been evaluated, in TBEV infected brain sections, very few CD4 T cells were detected and CD8 T cells were suggested to mediate immunopathology(^71, 72^). We evaluated T cell infiltrates by immunostaining for CD4 and CD8 in POWV infected brain sections 15 dpi. Despite overt spongiform damage in the CNS, we found very few infiltrating CD4 T cells in any areas of the CNS in 10 or 50 week old mice, with areas of the pons and cerebellum depicting maximal CD4 staining (Figure S4A,B). The CD8 positive T cells observed were sparse and dispersed sporadically in the pons and cerebellum of 10 and 50 week old mice 15 dpi, with maximal CD8 positive areas of the CNS presented, and no quantifiable differences in 10 or 50 week old POWV infected brains (Figure S4A,B). These CD4 and CD8 T cell findings differ dramatically from the ubiquitous staining of microglia/macrophage and astrocytes within the CNS of 10 and 50 week old mice 15 dpi, and foster roles for phagocytic glial cells and their responses in lethal POWV encephalitis(^43, 44, 47, 58, 59, 73–78^).

### Age-dependent Lethality Is Linked to POWV CNS Load

Viral load in the CNS of LI9 infected mice was measured 5, 10, 15 and 30 dpi by qRT-PCR and compared to age-matched controls. LI9 RNA was detected in murine brains of all age groups 5-15 dpi (n=4), with the highest viral burdens in 30-50 week old mice 10 dpi (10^8^-10^10^ copies/gm) (Figure 6A). However, by 15 dpi, viral RNA in the brains of 10, 20 and 30 week mice decreased to ≤10^4^ copies/gm, with RNA levels in 40 week old mice decreased by 2-3 logs 15 dpi (Figure 6A). In contrast, LI9 RNA levels persisted in the brains of 50 week old mice increasing by 1-2 logs 15 dpi (Figure 6A). In surviving mice 30 dpi we found that POWV RNA levels were reduced in all aged mice to levels at or just above the level of detection (Figure 6A). RNAscope in situ hybridization (ISH) of positive-stranded POWV RNA in the CNS(^27, 79, 80^) revealed sporadic POWV RNA in single cell foci dispersed throughout the brain (Figure 6B,S5). Coincident with qRT-PCR data of whole brains, POWV RNA ISH staining was notably increased in the cerebral cortex and midbrain of 50 week old brain sections 15 dpi (Figure 6B,S5). Collectively, these findings demonstrate that irrespective of murine age, POWV LI9 is neuroinvasive, with maximal viral RNA loads in the CNS of 10-40 week old mice 10 dpi. At 15 dpi there is a dramatic decrease in POWV load in younger mice and a contrasting increase in POWV RNA in the CNS of 50 week old mice.

**Figure 6.**
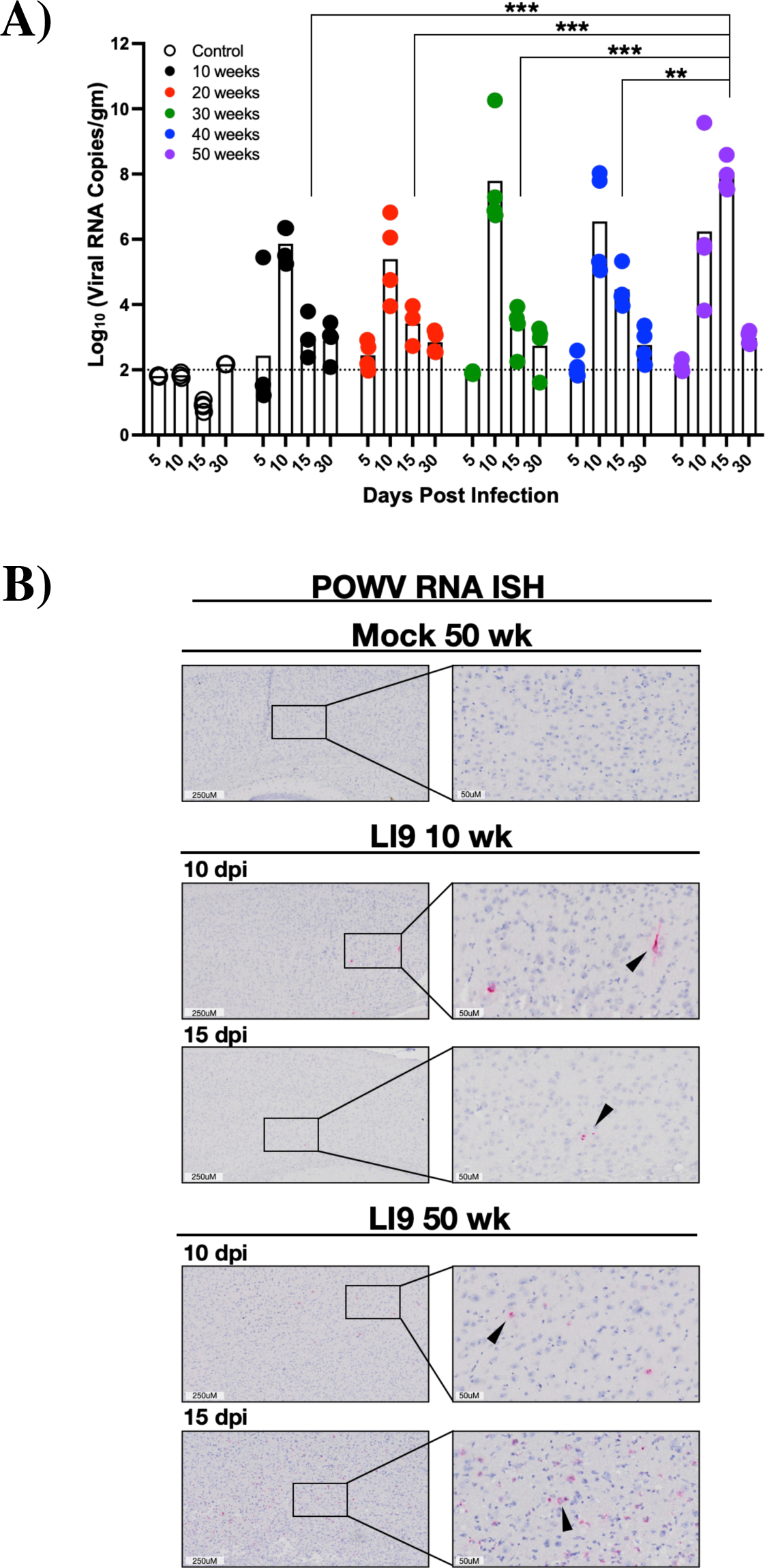
POWV CNS Load Kinetics and ISH Analysis. C57BL/6 10-50 week old mice were footpad inoculated with 2 × 10^3^ FFU of POWV LI9 or PBS. (A) POWV RNA levels in the CNS were assayed by qRTPCR 5-30 dpi and compared to mock infected controls. Data is expressed as the Log_10_ of POWV RNA (copies/gm) normalized to a standard curve. Each time point and age group reflect an n=4 following POWV infection, and n=11 for mock infected controls. Comparisons were performed by 2-way ANOVA analysis of log_10_ transformed values. Asterisks indicate statistical significance (**p*<*0.001, ***p*<*0.0001). (B) Brains from POWV infected or mock infected 10 and 50 week old mice were harvested 15 dpi, and genomic POWV RNA in the CNS was detected by in situ hybridization (ISH) with POWV RNAscope probe-red. Representative images of POWV RNA ISH in the cerebral cortex of POWV infected (n=3) and mock infected 50 week old mice (n=1) are presented. Black arrows indicate representative cells positive for genomic POWV RNA by ISH.

### POWV Directs Age-dependent Cytokine Responses that Distinguish CNS Microglial Phenotypes

Activated glia cell responses promote T helper (Th) cell effector functions by directing pro-inflammatory Th1-type cytokines or anti-inflammatory Th2 cytokine responses that in the CNS opposingly contribute to neuropathology or neuroprotection(^1, 44, 46, 52, 57, 59, 62, 63, 78^). We assayed the CNS of POWV LI9 infected 10 and 50 week old mice (n=3) for induced cytokine and chemokine responses 15 dpi by qRT-PCR and compared responses 5, 10 and 15 dpi to age-matched controls (Figure 7A). A comparison of CNS cytokines induced by POWV infection included a dramatic 5-154 fold induction of pro-inflammatory Th1-type cytokines (IL-2, IL-6, IL-12, IL-15, TNFα, IL-1β, IFNγ) in 50 week old mice over 10 week old mice 15 dpi (Figure 7B, S6)(^50–52, 55, 81, 82^). In contrast, in 10 week old mice POWV induced anti-inflammatory Th2-type cytokines IL-10 and TGF-β (^53, 57, 59, 61, 63, 83, 84^), that were 3-5 times lower in 50 week old mice (Figure 7B). IL-4 was induced by POWV in both 10 and 50 week old brains and this cytokine directs both Th2-type responses and, in combination with IL-2 or IL-12, inflammatory Th1-type TNFα and IFNγ responses(^51, 56, 59, 81^). Chemokines CCL2, CCL5 and CXCL10 were induced in both 10 and 50 week old mice, however induction was 3-5 times higher in 10 week old mice (Figure 7B). Kinetic analysis revealed that IL-2, IL-12, TNFα and IFNγ were only induced 15 dpi in 50 week old mice, while IL-10, TGFβ, SOCS3, CCL2, CCL5 and CXCL10 were induced 3-5 fold higher in 10 versus 50 week old mice, 15 dpi (Figure 7A,B,S6). CNS responses in 50 week old mice are consistent with inflammatory M1 microglial activation directing neurodegenerative Th1-type cytokines (Figure 7B, red/white), while in 10 week old mice distinct neuroprotective Th2-type cytokine responses elicited (Figure 7B, blue) are consistent with M2 microglial activation(^57, 59, 63, 85, 86^). These findings reveal a mechanism of age-dependent POWV lethality resulting from POWV neuroinvasion, increased CNS viral load, and the differential induction of late, inflammatory versus neuroprotective, cytokine responses. In addition, these results suggest potential therapeutic targets for resolving POWV lethality in a murine model that mirrors POWV disease and persistent spongiform encephalopathy in the elderly(^16^).

**Figure 7.**
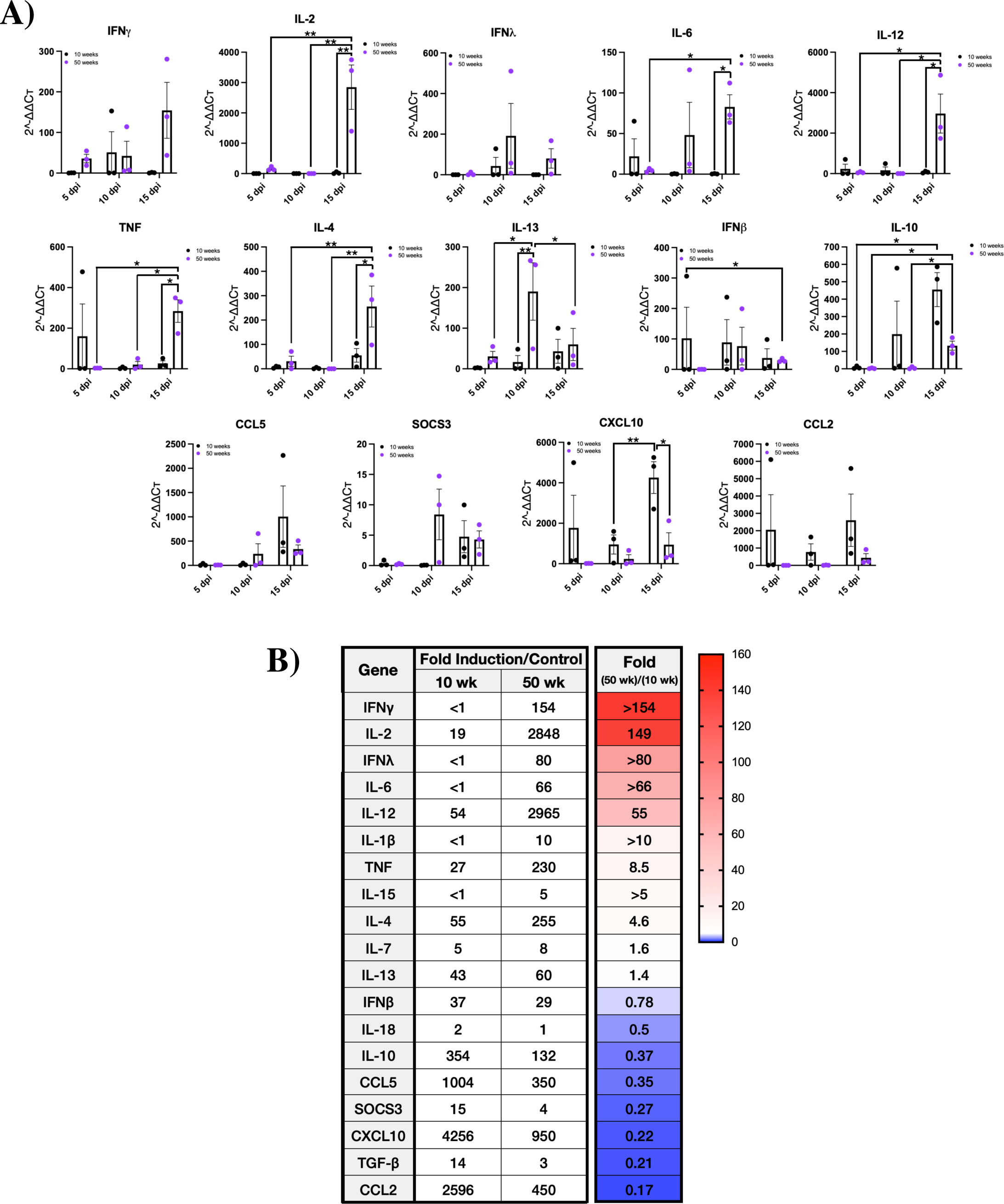
POWV Induction of CNS Cytokines and Chemokines is Age-dependent. (A) C57BL/6 10 and 50 week old mice were footpad inoculated with 2 × 10^3^ FFU of POWV LI9 or mock infected with PBS. Brains from POWV or mock infected mice were harvested, total RNA extracted and the induction of cytokine and chemokine transcripts were analyzed kinetically (5, 10 and 15 dpi) for fold induction over aged matched controls by qRT-PCR after standardizing to GAPDH RNA levels. Each time point, age group and infected or mock infected age-matched control is presented (n=3). Comparisons to mock infected controls were performed by one-way ANOVA analysis. Asterisks indicate statistical significance (\**p<*0.05; \*\**p<*0.01) (B) A summary of the fold induction of 10 and 50 week old CNS cytokine and chemokine responses over age-matched controls 15 dpi is presented. Findings are sorted by the fold induction of 50 week/10 week old CNS responses with fold differences and a heat map that segregates age-dependent responses.

## Discussion

Severe encephalitis and long-term neurologic damage are associated with human POWV infections(^2, 16^), yet mechanisms of POWV neuropathology remain largely unknown. Despite a lack of systemic infection or peripheral disease, POWV patients display profound cerebellar and brainstem involvement, characterized by neuronal loss, microgliosis, inflammatory infiltrates, and POWV RNA in the CNS(^2, 16^). Although human POWV cases are limited, available data indicates that lethal POWV infections occur in individuals >60 years of age(^2, 7, 14, 17^). This is consistent with the age-dependent lethality of TBEV infections(^38, 39^) and the severity of mosquito-borne West Nile virus (WNV) and Japanese Encephalitis virus (JEV) infections(^73, 87–89^).

Models of POWV infection were previously studied in peripherally inoculated 5-14 week old mice using murine brain neuroadapted LB and SP strains(^2, 12, 24, 25, 27, 29, 34^). In these studies, lethality varied by strain, dose, kinetics, and suggested requirements for tick saliva in directing neurovirulence(^29, 33^). Our analysis was performed using mice 10-50 weeks of age and POWV LI9, a strain from circulating *Ixodes* ticks isolated in VeroE6 cells(^21, 42^). Footpad inoculation of POWV LI9 into 10-50 week old C57BL/6 mice resulted in 82% lethality in 50 week old mice, with lethality sequentially reduced with age to 7.1% in 10 week old mice (Figure 1B,E). LI9 lethality was dose independent and conferred by a minimal infectious dose (Figure 1D,E). These findings establish the age-dependent lethality of POWV LI9 in a murine model and reflect severe human POWV disease in the elderly(^13^).

Similar to CNS degeneration observed in POWV patients, histopathology of POWV LI9 infected mice revealed spongiform CNS pathology lesions in cerebellar and hindbrain regions of the pons, medulla and brainstem that were accompanied by neuronal depletion and microglial nodules(^2, 13, 16^). In contrast to prior studies, Purkinje cell neurodegeneration was not observed in murine CNS sections following POWV LI9 infection. However, intense staining of microglia/macrophages and GFAP positive astrocytes in the murine CNS 5-30 dpi denotes inflammatory activation that is consistent with the ongoing CNS inflammation and microglial involvement in encephalitic POWV patients.

POWV RNA was detected in murine brains from all age groups with the highest viral load in the CNS observed 10 dpi in 10-40 week old mice. In 10-40 week old mice, POWV RNA in the CNS was reduced by 2-6 logs between 10 and 15 dpi. Coincident with LI9 lethality, in 50 week old mice, POWV RNA loads were not reduced between 10 and 15 dpi and reached their highest level 15 dpi. Collectively, these findings are consistent with reduced lethality resulting from POWV clearance from the CNS in young mice, and increased POWV load enhancing lethality in 50 week old mice. In surviving mice 30 dpi, we observed POWV RNA in the CNS reduced to levels at the limit of detection.

Despite age-dependent lethality, LI9 is fatal in a fraction of younger mice, demonstrating that POWV neuroinvasion is age-independent. Spongiform damage to the CNS was observed in all mice by 5 dpi indicating early CNS entry and neurovirulence. These findings suggest that POWV LI9 causes biphasic CNS pathology that results in early CNS damage (5 dpi) independent of murine age, and lethality 10-15 dpi that is age-dependent. The cause of early neuronal loss and spongiform damage following POWV infection remains a key question fundamental to neurodegenerative and cognitive deficits in survivors of POWV infections irrespective of age(^2, 7, 9, 14, 16, 17, 90^).

Comparing the CNS of POWV infected 10-50 week old mice revealed common inflammatory histopathology 5, 15 and 30 dpi. In the CNS of all murine age groups, we observed increases in Iba1 positive microglia/macrophages and GFAP positive astrocytes that reflect activated inflammatory responses of CNS resident glial cells(^44–49, 58, 59, 62, 64, 65, 85, 91, 92^). Enhanced glial cell staining was marginally increased 5 dpi, but dramatically increased by 15 dpi, and observed throughout the CNS of all age groups. In addition, increased microglia/macrophage staining was pervasive in the CNS of surviving mice 30 dpi. These findings mirror TBEV directed activation of glial cells in murine and human brains(^93, 94^) and are consistent with a persistent glial cell inflammatory state in the CNS contributing to the long-term neurologic sequalae observed in POWV patients.

Activated glial cells are principal mediators of neuroinflammation and age-dependent neurodegenerative diseases(^44, 58, 59, 63, 65, 69, 85, 95^). Microglia are resident CNS immune cells that mediate innate phagocytic responses and subsequent repair by distinct neuroprotective or neurotoxic responses(^44, 58, 59, 62, 63, 85^). In a simplified view, resting microglia are activated into pro-inflammatory M1 or, neuroprotective M2 microglial phenotypes, that likely form a continuum of intermediate types dependent on localized cytokine and cellular stimuli(^59, 85, 86^). M1 microglia are activated by IFNγ and highly secrete proinflammatory cytokines (TNFα, IFNγ, IL-1β, IL-2, IL-6, IL-12) and immune cell recruiting chemokines(^44, 58, 59, 63, 65, 69, 85, 95^). In contrast, CCL2 activated M2 microglia secrete immunoregulatory (IL-10, TGFβ) and neuroprotective (IL-4, IL-13) cytokines. However, IL-4 in the context of IL-2 and IL-12 reportedly enhances pro-inflammatory Th1 responses by potentiating IL-12 induced proliferation, STAT3 activation and IFNγ secretion(^51, 56, 57^). Astrocytes similarly maintain CNS homeostasis, but on activation induce GFAP, glial scar formation, S100B activation of microglial RAGE receptors and IL-12 directed inflammatory responses that augment microglial inflammation in the CNS(^46–48, 56, 69, 77, 81, 96^). As a result, altering the balance of neurotoxic and neuroprotective glial cells in the CNS may direct damaging inflammatory CNS hyperactivation during POWV infection of the CNS(^44, 47, 59, 62, 63, 69, 74, 76–78, 85, 97^).

The activation of glial cells in all POWV infected mice suggests that divergent age-dependent CNS responses may determine POWV clearance and lethality(^44, 74, 76, 78, 97, 98^). We compared CNS cytokine and chemokine responses to POWV infection and found that in 10, but not 50 week old mice POWV induced 1000-4200 fold increases in chemokines (CCL2, CCL5, CXCL10) along with immunosuppressive IL-10 and TGFβ (354-, 14-fold) 15 dpi. Contrastingly, in POWV infected 50 week old mice we found highly induced pro-inflammatory cytokines IL-2 and IL-12 (>2800-fold) and TNFα, IL-6, IFNγ and IL-1β (230-, 66-, 154-, 10-fold) in the CNS. IL-4 was induced 5-fold higher in 50 versus 10 week old mice, but in 50 week old mice IL-4 induction is accompanied by the robust induction of inflammatory IL-2 and IL-12 cytokines, suggesting a proinflammatory IL-4 role in this context(^51, 56, 59, 81, 85, 99^).

The collaborative high level induction of IL-2, IL-6, IL-12, TNFα, IL-1β, IL-4 and IFNγ in the CNS of LI9 infected 50 week old mice reveals a neurodegenerative M1 microglial, Th1-type cytokine profile, that may contribute to CNS damage and POWV lethality(^51, 52, 55, 56, 59, 63, 81^). In contrast, the induction of IL-10, TGFβ, IL-4 and chemokines in the CNS of 10 week old mice reflects a neuroprotective, anti-inflammatory M2 microglial, Th2 response(^57, 59–61, 63^), supporting POWV clearance from the CNS. Thus ubiquitous glial cell activation results in age specific M1/Th1 or M2/Th2 type cytokine responses as neuroinflammatory determinants(^78, 81^) of age-dependent POWV lethality. The high level induction of Th1 cytokines in POWV infected 50 week old mice suggests roles for activated glial responses in failed viral clearance from the CNS, age-dependent POWV lethality and persistent neuroinflammation. The CNS transcriptional responses observed during POWV infection are limited by whole brain analysis where the source and effector targets of secreted cytokines are intermingled. These initial findings set the stage for single-cell RNA sequencing approaches to kinetically define CNS cell responses of aged versus young mice that govern POWV survival and lethality.

Glial cells are activated in TBEV, WNV and JEV encephalitis, but glial activation was not evaluated for directing discrete CNS cytokine responses that contribute to age-dependent severity(^40, 73, 75, 89, 95^). In WNV encephalitis, suggested increases in CD4/CD8 T cells and CD8 reactivation are reported to distinguish clearance in young versus aged mice(^73, 87^). In contrast, we observed few CD4/CD8 T cell infiltrates in the CNS, and no discernible differences in T cell infiltrates in POWV infected 10 versus 50 week old mice(^73, 89^). Our findings associate glial cell activation with discrete CNS cytokine responses that direct a persistent neuroinflammatory state, age-dependent POWV lethality and long-term CNS pathology. Further, these results suggest potential therapeutic targets for preventing severe/lethal disease in aged mice and POWV infections of the elderly.

In a recent study, POWV SP intraperitoneally inoculated into 6 week old C57BL/6 mice resulted in a 17% fatality rate with surviving mice having persistent meningeal inflammation and SP viral RNA detected in the CNS 56-84 dpi(^27^). Our analysis shows that POWV LI9 RNA is cleared from the CNS to levels at or just above control levels 30 dpi, but with persistent glial cell inflammation throughout the CNS. Like SP, this may reflect LI9 persistence(^27^), or result from persistent auto-amplifying inflammatory CNS responses. Whether the use of very young mice or POWV SP(^27^), serially passaged in murine brains(^25^), account for differences in murine persistence may be revealed using POWV reverse genetics(^42^). In contrast to prior studies, our findings reflect age-dependent POWV lethality, CNS inflammation and long term spongiform damage observed in POWV infections of the elderly.

In the CNS, age-dependent increases in microglial senescence(^99–101^) and the expression of IFNγ, IL-2 and IL-12 are linked to neurodegenerative disease and amplified Th1 responses(^51, 52, 55, 56, 81, 102^). IL-12 and Th1 responses in the CNS are associated with experimental autoimmune encephalitis (EAE), multiple sclerosis and the age-dependent cognitive deficits in Alzheimer’s disease (AD) and Parkinson’s disease (PD)(^59, 78, 81, 103–107^). IL-12 expression and Th1 responses are inhibited by steroids, which enhance Th2 cytokine responses and blocking IL-12 function reduces AD pathology(^81^). There is a single report that 5 POWV patients >60 years of age treated with corticosteroids survived, while POWV was lethal in 5/5 untreated patients >60 years of age(^14^). Similarly, dexamethasone treating 14 WNV patients was suggested to reduce acute disease and enhance recovery. Although steroids may hinder viral clearance, these anecdotal findings suggest the potential utility of steroids in suppressing late neuroinflammatory damage, and rationalize analyzing steroids as potential therapeutics in our lethal POWV murine model.

Neuronal damage directs CNS tissue regeneration through neuronal stem cell (NSC) and microglial proliferation however, microglial activation can also inhibit CNS regeneration(^97, 108–110^). Aging is known to decrease regenerative NSCs and increase senescent microglia and T cells that secrete pro-inflammatory cytokines(^97, 100, 109, 111–113^). In AD and PD, age-dependent accumulation of senescent NSCs and microglia were associated with a severe proinflammatory state(^100, 111^). Cytokine responses in the CNS of POWV infected 50 week old mice reflect induced senescent cell cytokines (TNFα, IL6, IL-1β) that may contribute to an age-dependent senescence associated secretory phenotype (SASP)(^2, 13, 16, 103, 105, 114^). However, whether CNS stem cell repair, senescence or a neurotoxic state of inflammatory glial cell activation contribute to age-dependent POWV lethality remains to be examined.

Our findings reveal a novel age-dependent model of lethal POWV encephalitis and long-term neurologic damage in survivors. Histopathology of POWV infected aged murine brains mimics POWV induced CNS pathology in fatal human cases(^16^) and long-term spongiform damage observed in survivors. Our results establish an age-dependent model of lethal POWV encephalitis in mice, and reveal the direct involvement of glial cell M1 versus M2 cytokine responses in distinguishing POWV clearance and survival in young mice, from age-dependent lethality in aged 50 week old mice. Our studies are consistent with lethality resulting from persistently activated glial cells, a neurodegenerative M1 type cytokine response and failed POWV clearance from the CNS in aged mice. This foundational murine model sets the stage for defining protective responses of young mice and age-dependent changes that confer POWV lethality in 50 week old mice using single cell sequencing and murine knockout approaches. Further analysis of CNS responses in young versus aged mice, is likely to permit analysis of specific therapeutic targets and approaches for resolving age-dependent POWV lethality in elderly patients that may apply to other age-dependent viral and chronic neurodegenerative diseases.

## SUPPLEMENTAL FIGURES

**Figure S1. POWV Causes Neuronal Depletion Without Disrupting Purkinje Cells.**

(A-B) C57BL/6 10 and 50 week old mice were footpad inoculated with 2 × 10^3^ FFU of POWV LI9 or mock infected with PBS. (A) Brains were harvested 5 and 15 dpi, and immunostained for neurons (n=3) using NeuN antibody. Representative sections of the Pons are presented. (B) Brains were harvested 5, 15, or 30 dpi, sectioned, and H&E stained. Representative images of the cerebellum in POWV LI9 infected and mock infected 50 week old mice are presented. Black arrows indicate representative Purkinje Cells.

**Figure S2. Kinetic Changes in the Pons of POWV Infected 20, 30 and 40 Week Old Mice.** (A) C57BL/6 mice 20, 30 and 40 weeks of age were footpad inoculated with 2 × 10^3^ FFU of POWV LI9 or mock infected with PBS. Brains were harvested 5 and 15 dpi, and H&E stained (n=4), or immunostained for microglia/macrophages (n=3) using anti-Iba1 antibody. Representative sections of the Pons are presented. (B) The severity of POWV directed spongiform encephalopathy, microgliosis, and neuronal necrosis in H&E stained Pons regions (n=4) were scored on a scale of 0-4 by blinded comparison versus age-matched controls: (0) baseline determined by control brain staining in select region, (1) localized lesion, (2) multiple localized lesions, (3) lesions spread throughout most of select region, (4) lesions uniformly spread throughout select region. Comparisons to mock infected controls were performed by two-way ANOVA analysis. Asterisks indicate statistical significance (*, *P<*0.01; **, *P<*0.001, ***, *P<*0.0001).

**Figure S3. POWV Causes Spongiform Encephalitis, Microgliosis and Neuronal Necrosis in Mice of All Ages.** C57BL/6 mice 10-50 week old were footpad inoculated with 2 × 10^3^ FFU of POWV LI9 or mock infected with PBS. Brains were harvested 15 dpi, sectioned and H&E stained (n=4). Representative images of CNS histopathology in the Pons, Medulla, Brainstem, Cerebellum, Midbrain and Cerebral cortex of POWV versus mock infected 50 week old brains 15 dpi were scored for spongiform encephalopathy, microgliosis, and neuronal necrosis (n=4) were scored as in Figure S2. Comparisons to mock infected controls were performed by one-way ANOVA analysis. Asterisks indicate statistical significance (*, *P<*0.01; **, *P<*0.001, ***, *P<*0.0001).

**Figure S4. POWV Infected CNS Contains Few CD4 and CD8 T cell Infiltrates.**

(A-B) C57BL/6 10 and 50 week old mice were footpad inoculated with 2 × 10^3^ FFU of POWV LI9 or mock infected with PBS. Brains were harvested 15 dpi and immunostained for T cell infiltrates using CD4 or CD8 antibodies (HistoWiz). (A) Representative sections of the Pons of POWV infected 10 and 50 week old mice 15 dpi versus mock infected 50 week controls are presented. (B) Representative sections of the Cerebellum of POWV infected 10 and 50 week old mice 15 dpi versus mock infected 50 week controls are presented. Black arrows indicate representative CD4 or CD8 immunostained cells.

**Figure S5. POWV Infected CNS In Situ Hybridization.** C57BL/6 10-50 week old mice were footpad inoculated with 2 × 10^3^ FFU of POWV LI9 or PBS. (Brains from POWV infected or mock infected 10 and 50 week old mice were harvested 15 dpi, and genomic POWV RNA in the CNS was detected by in situ hybridization (ISH) with POWV RNAscope probe-red. (A) Pons; (B) Cerebellum; (C) Midbrain; (D) Medulla. Representative images of POWV RNA ISH in POWV infected (n=3) and mock infected 50 week old mice (n=1) are presented. Black arrows indicate representative cells positive for genomic POWV RNA by ISH.

**Figure S6. POWV Induced Responses of 50 vs 10 Week Old Mice, 15 dpi.** C57BL/6 10 and 50 week old mice were footpad inoculated with 2 × 10^3^ FFU of POWV LI9 or mock infected with PBS. Brains from POWV or mock infected mice were harvested, total RNA extracted and the induction of cytokine and chemokine transcripts were in the CNS were assayed by qRT-PCR, standardized to GAPDH RNA levels and compared to mock infected age-matched controls. Each time point, age group and infected or mock infected age-matched control is presented (n=3). Asterisks indicate statistical significance (\**p<*0.05; \*\**p<*0.01)

## DATA AVAILABILITY STATEMENT

The methods underlying this study are available an online supplement at the authors’ institutional website.

## ACKNOWLEDGEMENTS

We thank Jorge Benach for manuscript feedback and discussions of tick-borne diseases and neuropathogenesis; Brian Sheridan for discussion of CNS immune responses and cytokine regulation; Dan Salamongo for discussions of POWV disease and quantifying CNS responses; and Stephen Gaudino, for guidance on immunohistochemistry.

## MATERIALS AND METHODS

### Cells and Virus

VeroE6 cells (ATCC CRL 1586) were grown as previously described in Dulbecco’s modified Eagle’s medium at 37°C in 5% CO_2_. POWV strain LI9 (GenBank accession number: MZ576219) was isolated from infected *Ixodes scapularis* ticks via inoculation on Vero E6 cells, and stocks passaged 3-4 in VeroE6 cells as previously described(^21, 42, 115^). POWV LI9 was adsorbed onto 60% confluent VeroE6 monolayers for 1 h. Monolayers were PBS washed and grown in DMEM 5% FBS and POWV titers were determined by serial dilution and quantifying infected VeroE6 cell foci 1 dpi by immunoperoxidase staining with anti-POWV hyperimmune mouse ascites fluid (HMAF; 1:5,000 [ATCC]), as previously described(^21, 42, 60^). All the work with infectious LI9 POWV was performed in a certified BSL3 facility at Stony Brook University.

### Murine inoculation

C57BL/6J mice (10-50 weeks old) were purchased from Jackson Laboratory. Mice were anesthetized via intraperitoneal injection with 100 mg·ml^−1^ of ketamine and 20 mg·ml^−1^ of xylazine per kilogram of body weight. Animals were infected via subcutaneous footpad injection with up to 2 × 10^3^ FFU POWV or PBS in a volume of 20 μl. Mice were monitored daily for weight loss and signs of disease, and 5, 10, 15, 30 dpi mice were CO_2_ euthanized.

### Biosafety and Biosecurity

Animal research was performed in accordance with institutional guidelines using approved experimental protocols, and supervised by the Institutional Biosafety and Institutional Animal Care and Use Committees at SBU. Animals were managed by the SBU Division of Laboratory Animal Resources, which is accredited by the American Association for Accreditation of Laboratory Animal Care and DHHS, and maintained in accordance with the Animal Welfare Act and DHHS ‘Guide for the Care and Use of Laboratory Animals’. Veterinary care was directed by full-time resident veterinarians accredited by the American College of Laboratory Animal Medicine. Experiments with infectious POWV were performed in biosafety level 3 containment (SBU ABSL3 facility).

### Histopathological Analysis

Brains were harvested postmortem, fixed in neutral buffered formalin for 7 days, dehydrated with 70% ethanol for 24 h and paraffin embedded. Formalin-fixed paraffin-embedded (FFPE) brain tissues were sectioned (10 μm thickness) and histochemical staining with hematoxylin and eosin (H&E) was performed by the SBU Histology Core Lab. Immunohistochemical staining was performed by HistoWiz, Inc. with anti-Iba1 antibody (Wako 019-19741), GFAP (NB300-141), CD4 (ab183685), CD8 alpha (CST98941) or anti-NeuN antibody (ab177487) used to identify microglia/macrophage, astrocytes, CD4 T cells, CD8 T cells and neurons, respectively. Stained tissue sections were analyzed using QuPath software (https://qupath.github.io).

Histopathology scoring of H&E stained age-matched brain tissues was performed blinded to age and experimental group. Brain regions were scored on a scale of 0-4 for spongiform encephalopathy, microgliosis and neuronal necrosis. Scores define localized severity of pathology: (0) baseline of age-matched control brain staining in select region, (1) localized lesion, (2) multiple localized lesions, (3) lesions spread throughout most of select region, (4) lesions uniformly spread throughout select region. ImageJ quantification of Iba1 immunostaining of 50 week old mouse brains versus age matched mock infected controls: relative percent area of Iba1^+^ pixel intensity for 10 regions was determined per POWV infected brain and compared to mock infected controls.

For RNA in situ hybridization (ISH), FFPE brain tissues were sectioned (5 μm thickness), xylene deparaffinized, and underwent antigen retrieval in a decloaking chamber for 1 hr. POWV RNA was detected using the RNAScope Universal AP assay (Advanced Cell Diagnostics, Inc.) according to the manufacturer’s protocol. RNAScope 2.3 VS probe V-Powassan (catalog #415641) was used to detect genomic POWV RNA^(27)^. ISH probed slides were analyzed using QuPath software.

### RNA Extraction and qRT-PCR analyses

Age-matched mock or POWV-infected mice were euthanized and brains were harvested in TRIzol LS Reagent (Invitrogen) homogenized and RNA purified according to manufacturer’s protocols. RNA was processed using Monarch RNA Cleanup Kit (NEB T2030L) and quantified on a Nanodrop Spectrophotometer 2000. To define viral loads and inflammatory transcripts in POWV infected brains, qRT-PCR was performed on purified RNAs from brains of age-matched mock and POWV infected mice(^21, 42, 115^). cDNA synthesis was performed using random hexamer priming of a Transcriptor first-strand cDNA synthesis kit (Roche) as previously described (). Viral RNA levels were assayed using NS5 specific LI9 primers (Table 2) and compared to a standard curve of serially diluted POWV RNA. qRT-PCR transcript primers were designed using the NCBI gene database, with 60°C annealing profiles (Table 2). Transcript levels were analyzed in triplicate from n=3 age-matched control or n=3 POWV infected mice using PerfeCTa SYBR green SuperMix (Quanta Biosciences) on a Bio-Rad C1000 Touch system with a CFX96 optical module (Bio-Rad). Responses were normalized to internal GAPDH mRNA levels, and the fold induction was calculated using the threshold cycle (2^-ΔΔCT^) method for differences between age-matched mock and POWV infected RNA levels at each time point post-infection.

**Table 2.**
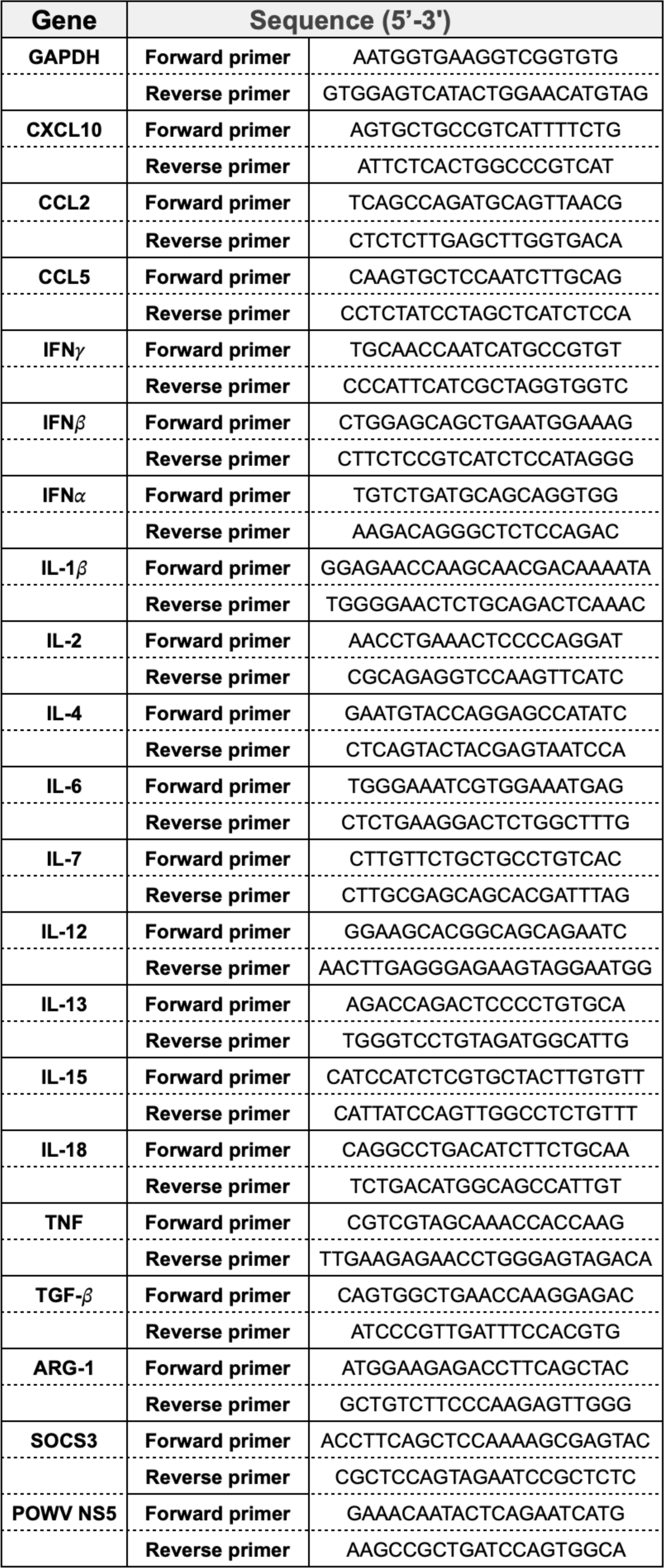
Primer Sequences Used for the qRT-PCR Detection of Select Transcripts.

### Neutralizing Antibody Assays

Neutralizing antibodies in sera from POWV LI9, or mock-infected mice were serially diluted and added to ∼500 FFU of POWV LI9 for 1 h. Sera-treated virus was adsorbed to 60% confluent VeroE6 cell monolayers for 1h and incubated. POWV infected cells were quantitated 36 hpi by immunoperoxidase staining with anti-POWV hyperimmune mouse ascites fluid (HMAF; 1:5,000 [ATCC]), horseradish peroxidase (HRP)-labeled anti-mouse IgG (1:2,000; KPL-074-1806), and 3-amino-9-ethylcarbazole (AEC) staining(^21,42,115^). CC50s were determined by reciprocal dilutions conferring a 50% reduction in infected cells versus controls.

### Statistical analysis

All of the details regarding the statistical analysis of the data can be found in the figure legends. The statistical significance of the results was determined by the use of Prism 6 software (GraphPad Software, Inc.; https://www.graphpad.com). Two-way comparisons were performed by two-tailed analysis of variance and an unpaired Student’s *t*-test or 2-way ANOVA. P values of <0.05, <0.01, <0.001 and <0.0001 were considered statistically significant as indicated.

